# Reciprocal sharing of extracellular proteases and extracellular matrix molecules facilitates *Bacillus subtilis* biofilm formation

**DOI:** 10.1101/2023.09.22.558988

**Authors:** Thibault Rosazza, Chris Earl, Lukas Eigentler, Fordyce Davidson, Nicola R Stanley-Wall

## Abstract

Extracellular proteases are a class of public good that support growth of *Bacillus subtilis* when nutrients are in a polymeric form. *B. subtilis* biofilm matrix molecules are another class of public good that are needed for biofilm formation and prone to exploitation. In this study, we investigated the role of extracellular proteases in *B. subtilis* biofilm formation and explored interactions between different public good producer strains across various conditions. We confirmed that extracellular proteases support biofilm formation even when glutamic acid provides a freely available nitrogen source. Removal of AprE from the NCIB 3610 secretome adversely affects colony biofilm architecture, while sole induction of WprA activity into an otherwise extracellular protease-free strain is sufficient to promote wrinkle development within the colony biofilm. We found that changing the nutrient source used to support growth affected *B. subtilis* biofilm structure, hydrophobicity, and architecture. We propose that the different phenotypes observed may be due to increased protease dependency for growth when a polymorphic protein presents the sole nitrogen source. We however cannot exclude that the phenotypic changes are due to alternative matrix molecules being made. Co-culture of biofilm matrix and extracellular protease mutants can rescue biofilm structure, yet reliance on extracellular proteases for growth influences population coexistence dynamics. Our findings highlight the intricate interplay between these two classes of public goods, providing insights into microbial social dynamics during biofilm formation across different ecological niches.

## Introduction

To persist in diverse environmental contexts, bacteria deploy various survival strategies, encompassing resistance to specific abiotic conditions, such as nutrient limitation (Harder and Dijkhuizen, 1983), extreme pressure (Merino et al., 2019), or elevated salinity (Oren, 2008). The survival strategies used also encompass interactions with biotic factors, including competitive dynamics, such as toxin secretion to impede the growth of neighbouring cells (Hibbing et al., 2010), or mutually beneficial collaborations, such as the division of labour with specific partners allowing coexistence with neighbouring species (West and Cooper, 2016).

In their native habitats, bacteria frequently form sessile, self-encased communities known as “biofilms” (Flemming et al., 2016). Within this context, a mechanism employed by bacteria to secure nutrients involves the synthesis and release of degradative enzymes that selectively target substrates, facilitating the breakdown of complex nutrient sources within their immediate surroundings (Hill et al., 2010, Armstrong, 2015, Cezairliyan and Ausubel, 2017). Illustrating this concept are chitinases, enzymes that break down chitin, a polymer composed of extended chains of N-acetylglucosamine (Hamid et al., 2013). Chitin is ubiquitous in various ecological systems, forming a structural component of fungal cell walls (Schonbichler et al., 2020) and serving as a fundamental element of insects and crustacean exoskeletons (Triunfo et al., 2022). The action of chitinases released by bacteria, such as those produced and secreted by *Vibrio cholerae*, enables the growth and the formation of cell clusters, as the resultant breakdown products serve as a carbon source (Drescher et al., 2014). The chitinases are a public good, as the digested chitin is available for all cells in the surrounding area, whether they contribute to the production of the enzymes or not (Drescher et al., 2014). Strategies, such as biofilm matrix hypersecretion leading to thick biofilms that retain enzymes locally (Drescher et al., 2014), have evolved to reduce sharing with non-producing cells thereby avoiding the so-called tragedy of the commons where the population collapses (Drescher et al., 2014).

*Bacillus subtilis*, a Gram-positive bacterium, has been the focus of extensive investigation concerning its ability to form robust, architecturally complex biofilms (for a review see (Arnaouteli et al., 2021)). The capacity of *B. subtilis* to develop biofilms has widespread ecological and industrial implications ranging from promoting plant health to healing fissures in concrete (Mahapatra et al., 2022, Mahmood et al., 2022). The transition from a free-swimming, planktonic state to a sessile, biofilm-associated mode of growth is governed by a complex network of genetic and environmental factors (Arnaouteli et al., 2021). Residents in the biofilm become encased in a self-produced extracellular matrix, which is primarily composed of polysaccharides, proteins, and nucleic acids (Flemming and Wingender, 2010). In the case of *B. subtilis*, the main biofilm matrix components include fibres formed by TasA (Branda et al., 2006), an exopolysaccharide (EPS) (Branda et al., 2001) that has an di*N*-acetyl bacillosamine-appended lipid anchor (Arbour et al., 2023), a film-forming protein called BslA (Kobayashi, 2007), and for some conditions and/or isolates, polyglutamic acid (Morris et al., 2022, Kalamara et al., 2021). These matrix molecules allow the attachment of *B. subtilis* to a surface (Flemming et al., 2007), allow the cells to adhere together (Branda et al., 2006), and provide other important biological functions including the formation of a hydrophobic layer on the surface of the biofilm (Kobayashi, 2007) to name but a few roles that are essential to support biofilm formation and promote the resilience of the resident cells.

*B. subtilis* has embedded regulatory mechanisms that drive the division of labour and the production of public goods to maximise biofilm productivity (Kearns, 2008, Dragos et al., 2018a, Martin et al., 2020). Well-documented examples of essential biofilm public goods are the extracellular matrix protein TasA and the EPS (Dragos et al., 2018a). The co-culture of TasA and EPS mono-producer strains increases the ability *B. subtilis* to form a biofilm at the air-liquid interface (Dragos et al., 2018a, Dragos et al., 2018b). This division of labour arises from the genetically distinct strains contributing to specific matrix components that surpass the wild-type strain producing all matrix components in a phenotypically heterogeneous manner This .In planktonic conditions, the eight extracellular proteases produced by *B. subtilis* serve as a public good that incurs a context-dependent cost that increases as the dependence on the extracellular protease activity for growth rises (Rosazza et al., 2023). The extracellular proteases have a wide array of functions which has been covered by a recent review (Harwood and Kikuchi, 2022). It has not been shown if the extracellular proteases function as a public good within biofilms. However, it is known that during biofilm formation, extracellular proteases are heterogeneously produced within the population (Marlow et al., 2014) and that the suite of eight secreted extracellular proteases is needed to form a wild-type-like colony biofilm, even when glutamic acid serves as the nitrogen source (Earl et al., 2020).

Here we explore the extent to which the extracellular proteases play a role in supporting biofilm formation by *B. subtilis*. We proposed that the extracellular proteases were a biofilm public good and that the extracellular proteases and matrix molecules could be shared for the mutual benefit of genetically defined non-producer strains during *B. subtilis* biofilm formation. Conventionally, the growth medium used for the analysis of *B. subtilis* biofilms contains a readily accessible and plentiful supply of glutamic acid as the nitrogen source. However, we proposed that these conditions could mask the role of the extracellular proteases in biofilm formation. As part of our study, the dependency upon extracellular proteases for growth and the impact on biofilm formation was explored by providing nitrogen in a polymeric form. We found that the extracellular proteases and matrix components could be shared to reinstate biofilm formation, but as the dependency on the extracellular proteases to sustain growth increased, the ability of the genetically defined non-producer strains to coexist in harmony was impacted and growth rather than biofilm formation is prioritised.

## Materials and Methods

### Strains and growth media used in this study

Strains used in this study were derived from *B. subtilis* isolate NCIB 3610 or NCIB 3610 *comIQ12L* (both are referred to as WT) (Konkol et al., 2013) or NCIB 3610 *comIQ12L ΔaprE, Δbpr, Δepr, Δmpr, ΔnprB, ΔnprE, Δvpr, ΔwprA* (referred to as Δ8 and stocked here as NRS5645) (Earl et al., 2019). See Table 1 for a full list of the genotypes. *B. subtilis* strains were routinely grown using Lysogeny-Broth (LB) medium (10 g NaCl, 5 g yeast extract, and 10 g tryptone per litre) at 37°C for 16 hours. For colony biofilm formation, *B. subtilis* strains were grown using an MS base medium (5 mM potassium phosphate and 100 mM MOPS at pH 7.0 supplemented with 2 mM MgCl_2_, 700 μM CaCl_2_, 50 μM MnCl_2_, 50 μM FeCl_3_, 1 μM ZnCl_2_, 2 μM thiamine) containing 0.5% (v/v) glycerol (abbreviated to Gly) and one of 0.5% (w/v) glutamic acid (abbreviated to Ga), 1% (w/v) BSA or 1.5% (w/v) milk as required and indicated. Where needed, the medium was solidified using 1.5% (w/v) Select Agar (Invitrogen). The growth medium containing glycerol and glutamic acid is frequently referred to as “MSgg” (Branda et al., 2001).

**Table 1.** Strains used in this study.

| Strain | Genotype<sup>a</sup> | Source/Construction<sup>b</sup> |
| --- | --- | --- |
| NCIB 3610 | Prototroph | *Bacillus* Genetic Stock Centre |
| DK420<br>(referred to here as NRS6017) | NCIB 3610 *comI*<sup>Q12L</sup> | (Konkol et al., 2013) |
| NRS5645 | NCIB 3610 *comI*<sup>Q12L</sup> $\Delta aprE$, $\Delta bpr$, $\Delta epr$, $\Delta mpr$, $\Delta nprB$, $\Delta nprE$, $\Delta vpr$, $\Delta wprA$ (referred to as $\Delta 8$) | (Earl et al., 2019) |
| NRS1473 | NCIB 3610 *sacA*-P*spachy-gfp* (*kan*) | (Verhamme et al., 2007) |
| TB35 (referred to as NRS6556) | NCIB 3610 *comI*<sup>Q12L</sup> *amyE::Phy-spank – mKate2* (*cml*) | Gift from Professor Akos T. Kovacs (Holscher et al., 2016) |
| NRS3648 | $\Delta 8$ *amyE::Phy-spank-gfp**mut2* (*cml*) | pBL165 (Stanley et al., 2003) → NRS5645 |
| NRS3670 | $\Delta 8$ *amyE::Phy-spank-mKate2* (*cml*) | pNW725 (Eigentler et al., 2022) → NRS5645 |
| NRS3657 | NCIB 3610 *comI*<sup>Q12L</sup> $\Delta aprE::erm$ | (Rosazza et al., 2023) |
| NRS6047 | NCIB 3610 *comI*<sup>Q12L</sup> $\Delta bpr$ | (Earl et al., 2019) |
| NRS3658 | NCIB 3610 *comI*<sup>Q12L</sup> $\Delta epr::erm$ | (Rosazza et al., 2023) |
| NRS3659 | NCIB 3610 *comI*<sup>Q12L</sup> $\Delta mpr::erm$ | (Rosazza et al., 2023) |
| NRS3660 | NCIB 3610 *comI*<sup>Q12L</sup> $\Delta nprB::erm$ | (Rosazza et al., 2023) |
| NRS3661 | NCIB 3610 *comI*<sup>Q12L</sup> $\Delta nprE::erm$ | (Rosazza et al., 2023) |
| NRS7010 | NCIB 3610 *comI*<sup>Q12L</sup> $\Delta vpr$ | (Earl et al., 2019) |
| NRS6810 | NCIB 3610 *comI*<sup>Q12L</sup> $\Delta wprA$ | (Earl et al., 2019) |
| NRS3679 | NCIB 3610 *comI*<sup>Q12L</sup> $\Delta 7 aprE^{WT}$ (*erm*) | (Rosazza et al., 2023) |
| NRS3680 | NCIB 3610 *comI*<sup>Q12L</sup> $\Delta 7 bpr^{WT}$ (*erm*) | (Rosazza et al., 2023) |
| NRS3681 | NCIB 3610 *comI*<sup>Q12L</sup> $\Delta 7 epr^{WT}$ (*erm*) | (Rosazza et al., 2023) |
| NRS3682 | NCIB 3610 *comI*<sup>Q12L</sup> $\Delta 7 mpr^{WT}$ (*erm*) | (Rosazza et al., 2023) |
| NRS3683 | NCIB 3610 *comI*<sup>Q12L</sup> $\Delta 7 nprB^{WT}$ (*erm*) | (Rosazza et al., 2023) |
| NRS3684 | NCIB 3610 *comI*<sup>Q12L</sup> $\Delta 7 nprE^{WT}$ (*erm*) | (Rosazza et al., 2023) |
| NRS3885 | NCIB 3610 *comI*<sup>Q12L</sup> $\Delta 7 sacA::kan vpr^{WT}$ | (Rosazza et al., 2023) |
| NRS6362 | NCIB 3610 *comI*<sup>Q12L</sup> $\Delta 7 wprA^{WT}$ | (Earl et al., 2019) |
| NRS3812 | NCIB 3610 *comI*<sup>Q12L</sup> $\Delta bsIA::cml$ *sacA::Phy-spank-gfp mut2* | (Hobley et al., 2013) |
| NRS3798 | NCIB 3610 *comI*<sup>Q12L</sup> $\Delta eps(A-O)::tet amyE::Phy-spank-gfp mut2$ | (Ostrowski et al., 2011) |
| NRS2984 | NCIB 3610 *comI*<sup>Q12L</sup> $\Delta tasA::(spc) amyE::Phy-spank-gfp mut2$ | (Ostrowski et al., 2011) |
| NRS6552 | NCIB 3610 *comI*<sup>Q12L</sup> $\Delta bsIA::cml$ $\Delta eps(A-O)::tet \Delta tasA::(spc) sacA::Phy-spank-gfp mut2$ | (Earl et al., 2019) |

<sup>a</sup>. Drug resistance cassettes are as follows: kan, kanamycin resistance; cml, chloramphenicol resistance; spec, spectinomycin resistance; tetracycline resistance, tet; and erm, erythromycin resistance.
<sup>b</sup>. Strain construction is denoted as DNA from donor strain transformed into the recipient strain following the direction of the arrow (→).

### Growing and imaging colony biofilms

Material from a −80°C glycerol stock was streaked onto an LB plate and incubated O/N at 30°C. A single colony was used to inoculate 5 mL of LB and incubated O/N at 200 rpm at 37°C. 100 µL of the O/N culture was added to 5 mL of LB and incubated at 200 rpm at 37°C. After 4 h, the culture was centrifuged for 10 min at 4500 rpm. The cell pellet was resuspended using 1 mL of base MS media (5 mM K_2_HPO_4_, 5 mM KH_2_PO_4_, 100 mM MOPS pH 7.0) and OD_600_ was measured. The culture density was normalised to an OD_600_ of 1 by adding base MS media as required. 10 µL was spotted to establish the colony biofilm. For the co-culture biofilms, the individual isolates were grown as described and mixed in 1 ml at a 1:1 ratio. 10µL of this mixed culture was used to establish the biofilm. The plates were incubated at 30°C for 1, 2, 3, 6 or 9 days. Depending on the type of experiment performed, biofilms were either imaged using a Nikon camera or a Leica M165C stereoscope or processed to measure biofilm hydrophobicity or extract their protein content. A separate plate was used for each biological replicate and for each time point. Images were uploaded and analysed using the OMERO or FIJI software.

### Measuring biofilm hydrophobicity

Biofilm hydrophobicity was analysed after 3 days of incubation at 30°C when the nitrogen source was either glutamic acid or milk and after 6 days for BSA as described previously (Hobley et al., 2013, Kalamara et al., 2021). A contact angle above 90° indicates that the biofilm is non-wetting and a contact angle value below 90° indicates that the biofilm is wetting.

### Quantifying biofilm footprint and the proportion of a fluorescent population within a colony biofilm

To quantify the biofilm footprint one of two methods was used. First, we used OMERO.Insight v.5.8.1 (Allan et al., 2012). Each biofilm was highlighted as a region of interest and the area it occupied was highlighted and calculated. The total area of the field of view and the theoretical maximum possible for a perfectly circular biofilm in the field of view are shown on the graphs. Note that for some colony biofilms, the area occupied will be larger than that reported as the edges of the structure extend beyond the field of view. To quantify the proportion of each fluorescent population inside the biofilm footprint, the total signal in each channel was derived for the region of interest (e.g., GFP and mKate2). The mean background signal per pixel (determined from images taken of a strain lacking the respective fluorescent protein under the same conditions) was subtracted considering the area occupied by the sample. The percentage of matrix mutant in the biomass was determined by calculating the total signal minus background in the FITC channel for the total signal minus background in both fluorescence channels. This method for quantifying the presence of two strains in a colony biofilm has been extensively validated in our previous work (Eigentler et al., 2022). Additionally, for Figure S3, the footprint occupied by the colony biofilms was quantified using a Python script using the Scikit-image plugin (van der Walt et al., 2014). Images obtained from the Nikon camera (.JPEG files) or the Leica M165C stereoscope (.LIF files) were imported into Python via the pillow or the read LIF plugin. In brief, the biofilm images were cropped and centred to retain the biofilm and remove the border of the Petri dish if it was present in the image. A multi-Otsu threshold was calculated to remove background and noise. The biofilm contour was determined using a “marching squares” method (Lorensen and Cline, 1987) to find iso-valued contours in a 2D array and the scale was fixed using a reference to yield the area.

### Protein extraction and Immunoblotting

For samples grown using glutamic acid 0.5% (w/v) or milk 1.5% (w/v) proteins were extracted after 3 days of incubation at 30°C. For samples grown using BSA 1% (w/v) proteins were extracted after 6 days of incubation at 30°C. A tablet of protease inhibitor (Roche) was dissolved into 5 mL of BugBuster™ protein extraction reagent (Sigma). Using a sterile loop, biofilms were gently recovered and then resuspended into an Eppendorf tube containing 0.5 mL of protease inhibitor BugBuster™ solution. Biofilms were disrupted by passing through a 23G needle fitted on a 1 mL syringe 5 times and then sonicated for 5 sec (2 times) at 30% amplitude with 10 sec intervals (“off time”). Tubes were incubated for 20 min at 21°C and then centrifuged for 10 min at 13 000 g at 4°C. A volume of 400 µL of supernatant was transferred into a new Eppendorf tube and protein extracts were stored at −20°C. To determine the concentration of protein a BCA assay kit was used (Pierce).

10 µg of protein mixed with 7.5 µL of 4X loading dye (0.5 M Tris-HCl pH 6.8, 0.1 M EDTA, 15.5% (w/v) SDS, 3% (v/v) glycerol, 5% (v/v) β-mercaptoethanol, 1 mg bromophenol blue) and H_2_O was added to bring the total volume to 30 µL. To denature proteins, tubes were incubated for 5 min at 95°C and centrifuged 1 min at 13 000 rpm. 30 µL samples were loaded onto a 10% (w/v) SDS-PAGE gel and run for ±1 h at 120 volts prior to transfer to a polyvinylidene difluoride membrane. The membrane was transferred to a 50 mL Falcon tube and washed using 1x TBS containing Tween 0.2% (v/v) and then incubated at 4°C on a roller shaker for 1 h in 5 mL of TBS Tween 0.2% (v/v) + milk 3% (w/v) solution. The membranes were washed and incubated O/N at 4°C on a roller shaker in 5 mL of TBS Tween 0.2% (v/v) + milk 3% (w/v) solution containing the primary antibody. For αBslA (rabbit) a 1:500 dilution was used and αTasA (rabbit) a 1:25,000 dilution was used (Ostrowski et al., 2011). Membranes were transferred into a new 50 mL Falcon tube and washed as described above prior to incubation in 5 mL of TBS Tween 0.2% (v/v) + milk 3% (w/v) containing αRabbit-HRP (goat) secondary antibody at a 1:5000 dilution. The membranes were washed as above and a mix of 1 mL ECL1 and 1 mL ECL2 solution (Thermo Fisher) was used for signal detection via an Azure scanner. Images were visualised using FIJI software.

### Determining biofilm productivity and sporulation percentage

After 3 days of incubation at 30°C, biofilms were transferred from the agar surface into 1mL of PBS 1X using a sterile loop. The biomass was disrupted by multiple passages through a 23G needle and sonication for 3 sec (2 times) at 20% amplitude with 10-sec intervals (“off time”). To determine the number of CFU/biofilm, 10-fold serial dilutions were performed up to 10^−8^ in 1 mL of PBS 1X and 100 µL of the 10^−8^ dilution was plated onto the surface of an LB agar plate. To determine the proportion of spores per biofilm, the remaining 900 µL of the 10^−8^ dilution was heat-treated for 20 min at 80°C and left to cool for 10 min at room temperature. 100 µL of this heat-treated sample was plated onto an LB agar plate after mixing. The agar plates were incubated O/N at 37°C and the number of CFUs on each plate was enumerated to calculate the CFU/biofilm and CFU spores/biofilm.

### Data analysis and visualisation

The datasets and code generated are available in a Zenodo repository (Rosazza et al., 2024). Data was plotted using Graphpad Prism version 9.4.1 and analysed using the built-in methods as specified.

## Results

### The extracellular proteases are needed for wild-type colony biofilm formation

NCIB 3610 (a wild-type (WT) protease-producing isolate) forms a rugose colony biofilm when glutamic acid is the main nitrogen source (Figure 1A, S1) (Branda et al., 2001). In the same conditions, the Δ8 strain (NRS5645 - a derivative of NCIB 3610 that lacks the coding regions of the eight known extracellular proteases (Earl et al., 2020)) forms an expanded colony biofilm where the interior lacks wrinkles and the periphery develops limited wrinkles after 3-days of incubation (Figure 1A, S1). The Δ8 strain also forms less architecturally complex pellicle biofilms (Earl et al., 2020). Quantification of the surface area occupied by the colony biofilms revealed a larger footprint for the Δ8 strain compared to the protease-producing parental strain at all time points from 48 hours onwards (Figure 1B). These results confirm and extend our previous work (Earl et al., 2020), demonstrating that when nitrogen is in the form of glutamic acid, the extracellular proteases support biofilm architecture development and restrict biofilm expansion.

**Figure 1.**
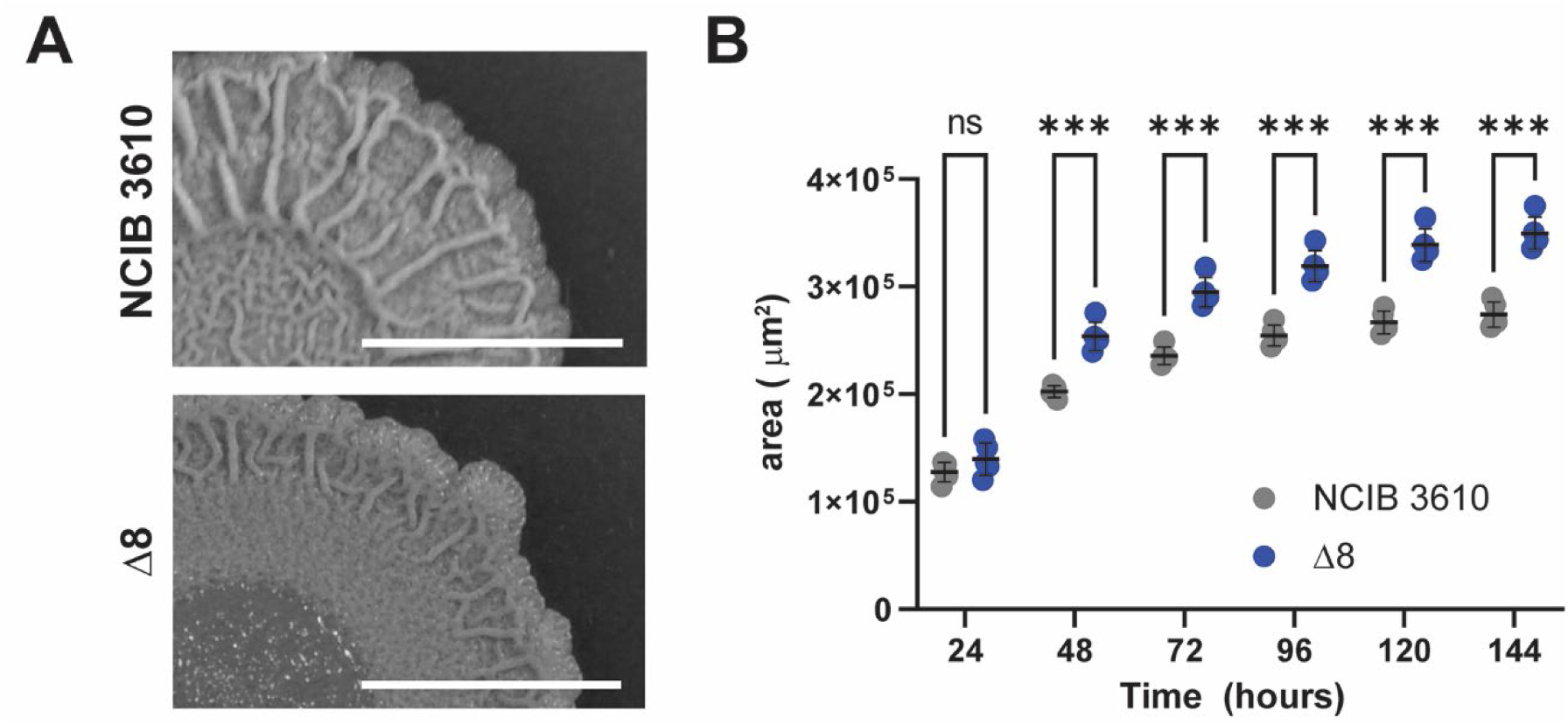
Extracellular proteases support the formation of a rugose colony biofilm. **A.** NCIB 3610 and Δ8 colony biofilms after 144 hrs incubation at 30°C (also shown in Figure S1). The scale bar represents 5 mm; B. Biofilm footprint area in μm^2^ of NCIB 3610 and Δ8 strains grown using 0.5% glutamic acid (w/v). Each point represents the footprint area value (n=3 biological replicates), the line represents the mean value, and the error bars are the standard deviation. Unpaired T-tests were performed between NCIB 3610 and Δ8 strain at each time point: *** = p-value < 0.001.

### The proteinaceous extracellular matrix components are produced in the absence of the extracellular proteases

We hypothesized that the Δ8 strain colony biofilm defect may be a consequence of it lacking one of the proteinaceous extracellular matrix components (namely BslA (Kobayashi, 2007) and/or TasA (Branda et al., 2006)). We assessed the ability of the Δ8 strain to produce BslA and TasA and used a triple extracellular matrix component mutant strain (Δ*bslA*, Δ*eps(A-O)*, Δ*tasA* (NRS6552) as a negative control for the immunoblot analysis. We grew colony biofilms for three days (Figure 2A) and extracted proteins from the entire biomass. Neither TasA nor BslA were detected in the negative control sample (Figure 2B). We detected both BslA and TasA proteins in the Δ8 biofilm samples (Figure 2B). In the NCIB 3610 protein extract, the TasA protein was present at its native molecular weight (∼28 kDa) and a lower molecular weight (∼10 kDa). In the Δ8 biofilm protein extract, the TasA protein was only present at its apparent native molecular weight (∼28 kDa). These results indicate that the two main extracellular matrix proteins are produced in the absence of extracellular protease activity. However, not producing the extracellular proteases impedes the cleavage of TasA and we have not assessed if the profile of transcription is impacted. The lack of TasA cleavage in the Δ8 strain may be responsible for the impact observed on colony biofilm formation, but this remains to be investigated.

### The absence of extracellular proteases increases the percentage of spores

We next tested if the phenotypic difference observed between NCIB 3610 and Δ8 strains during biofilm formation was due to a growth difference that emerged under sessile conditions. The growth difference could arise due to spatial constraints imposed during biofilm formation that physically force the cells to grow on top of one another (Asally et al., 2012, Fei et al., 2020) and that cells lyse (Asally et al., 2012), presumably releasing complex molecules into the biomass that would require extracellular proteases to access the nutrients. As a measure of growth, and thus biofilm productivity, we quantified the colony forming units per biofilm for the NCIB 3610, Δ8, and Δ*bslA* Δ*eps(A-O)* Δ*tasA* strains. As expected, the colony biofilm formed by the Δ*bslA* Δ*eps(A-O)* Δ*tasA* strain contained ∼700-fold fewer CFU/biofilm compared to NCIB 3610 (Figure 3A), confirming previous results showing that not producing biofilm matrix proteins impacts biofilm productivity (Dragos et al., 2018a). In contrast, there was no statistically significant difference in the number of colony-forming units per biofilm quantified between the NCIB 3610 and the Δ8 strain (Figure 3A).

**Figure 2.**
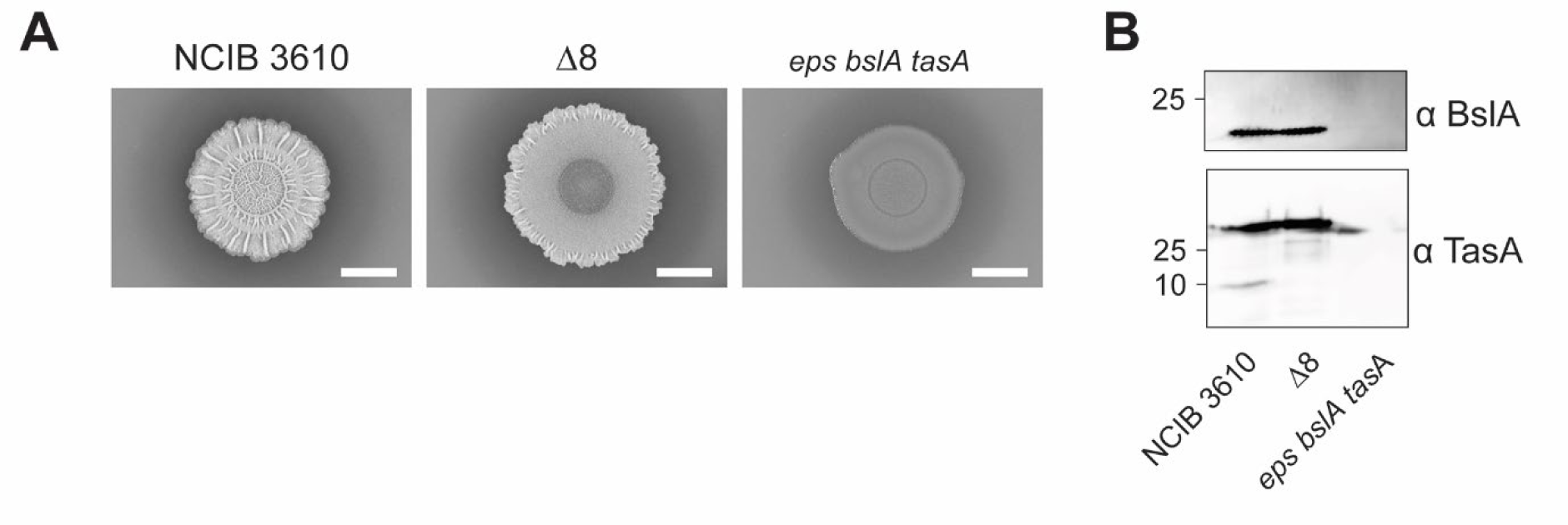
Extracellular matrix proteins are produced in the absence of extracellular proteases. **A.** NCIB 3610, Δ8 and Δ*bslA*, Δ*eps(A-O),* Δ*tasA* (*eps bslA tasA*) colony biofilms at day 3. A representative image of five independent experiments is shown. The scale bar represents 5 mm; **B.** Immunoblot analysis of biofilm protein extracts from NCIB 3610, Δ8 (NRS5645) and Δ*bslA*, Δ*eps(A-O),* Δ*tasA* (NRS6552) (labelled *eps bslA tasA*) probing BslA (top image) and TasA (bottom image) colony biofilms after 3 days. A representative image of three independent experiments is shown for the colony biofilms. All samples were grown using 0.5% glutamic acid (w/v) and 0.5% glycerol (v/v). The scale bar represents 5 mm.

**Figure 3.**
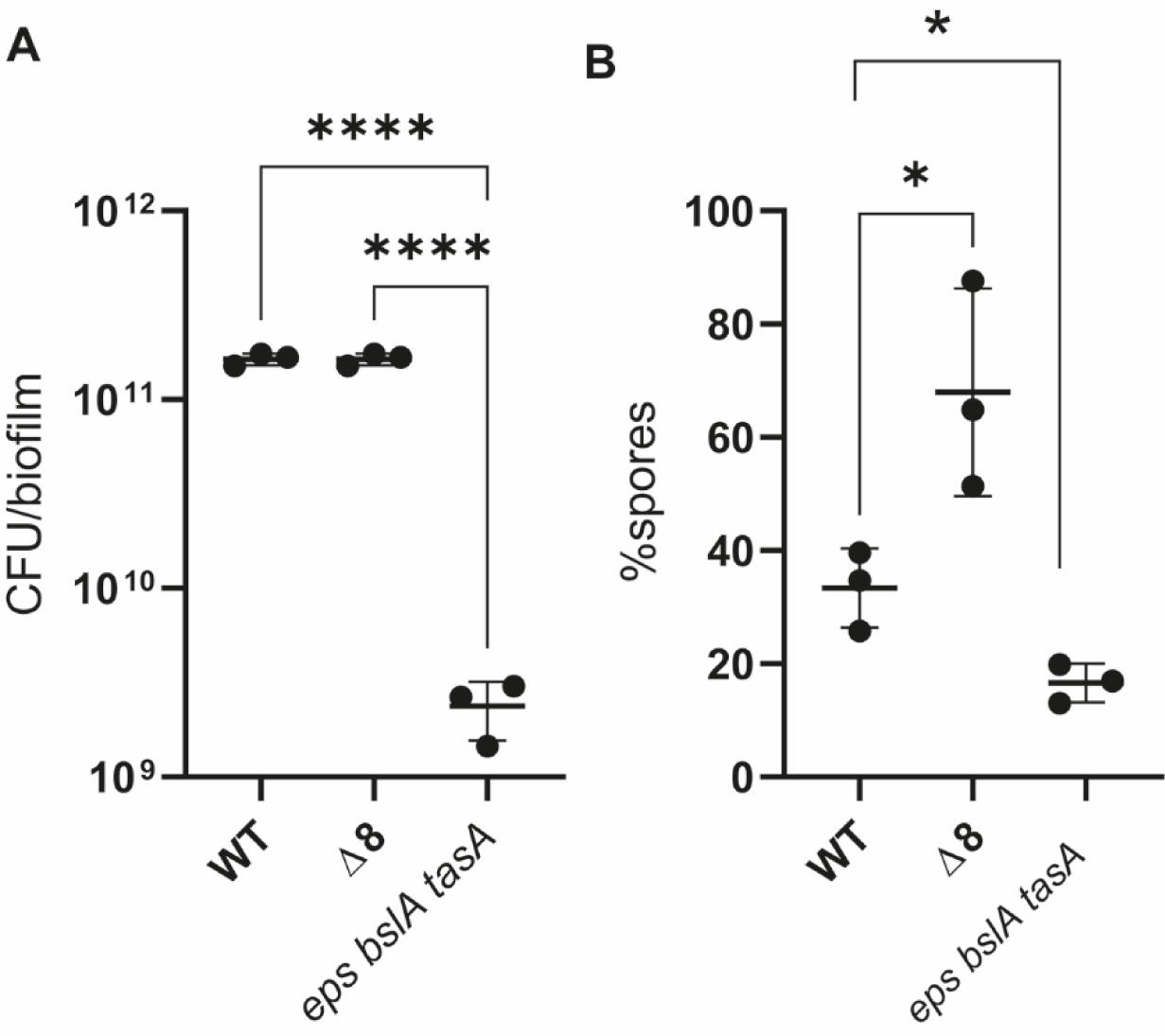
Sporulation within colony biofilms is impacted by the absence of extracellular proteases. **A.** Biofilm productivity at day 3 for NCIB 3610, Δ8 and a triple Δ*bslA*, Δ*eps(A-O),* Δ*tasA* triple deletion strain. Each point represents the CFU/biofilm (n=3 biological replicates) and the line represents the mean value. T-test performed between NCIB 3610 and Δ8 or triple deletion strains: **** = p-value < 0.0001; **B.** Proportion of spores per biofilm at day 3 for NCIB 3610, Δ8 and Δ*bslA*, Δ*eps(A-O),* Δ*tasA* strains. Each point represents the % sporulation (n=3 biological replicates) and the line represents the mean value. T-test performed between NCIB 3610 and Δ8 or triple deletion strains: * = p-value < 0.05. All samples were grown using 0.5% glutamic acid (w/v) and 0.5% glycerol (v/v).

Of the cells in the biofilm, the proportion of spores in Δ*bslA* Δ*eps(A-O)* Δ*tasA* triple mutant colony biofilms was lower (about half) than that of the NCIB 3610 biofilm (Figure 3B), confirming that not producing biofilm matrix proteins impedes the ability of the resident cells to sporulate (Vlamakis et al., 2008, Aguilar et al., 2010). The proportion of spores was consistently higher in Δ8 colony biofilms than in the wild-type strain (Figure 3B) with ∼2-fold (p<0.05) more spores in the protease-free strain. Our data show that the lack of extracellular proteases increases the frequency of cells sporulating during biofilm formation. The increase in spores may be linked with an inability to access nutrients in the founding part of the colony biofilm where local depletion of nutrients is possible (for example, (Arnaouteli et al., 2019)) and/or the impact of the anoxic environment that develops in the centre of the biomass (Flemming et al., 2016, Flemming and Wingender, 2010).

### The impact of specific extracellular proteases on colony biofilm formation

*B. subtilis* extracellular proteases have overlapping catalytic activities (Harwood and Kikuchi, 2022). We therefore predicted that there would be redundancy in the extracellular proteases required for wild-type colony biofilm formation. We postulated that the deletion of any single gene encoding an extracellular protease would have a limited impact on the colony biofilm phenotype. To explore this hypothesis, we used a suite of single extracellular protease mutant strains (see Table 1) (Rosazza et al., 2023). We spotted colony biofilms for each single gene deletion strain and imaged their formation over 3 days, a time frame where the phenotypic difference between NCIB 3610 and Δ8 becomes striking. We found that when *B. subtilis* does not produce AprE, the formation of wrinkles on the colony biofilm was impacted (Figure 4 and Figure S3). The *aprE* mutant colony biofilms were less structured, and the wrinkles were less abundant and prominent than those observed on NCIB 3610 colony biofilms. However, the phenotype of the *aprE* single mutant colony biofilm was different from that of the Δ8 strain with small wrinkles being placed across the entirety of the biofilm surface for the *aprE* mutant strain rather than solely at the extremity. In contrast, when *B. subtilis* did not produce one of Bpr, Epr, Mpr, NprB, NprE, Vpr or WprA there was no impact on the colony biofilm phenotype, with each single deletion strain exhibiting an NCIB 3610-like phenotype as has previously been shown for Vpr and NprB (Figure S3) (Earl et al., 2020, Dixit et al., 2002). In addition, we found that the area of the *wprA* mutant biofilm footprint was comparable to that of the parental NCIB 3610 strain at 72 hours (Figure S3). In contrast, the area occupied by the *epr* mutant biofilm at 72 hours was between that of the NCIB 3610 and Δ8 strain (Figure S3), while the remaining single deletion mutants had a similar biofilm footprint area to that of the Δ8 strain from 48 hours onwards (Figure S3).

**Figure 4.**
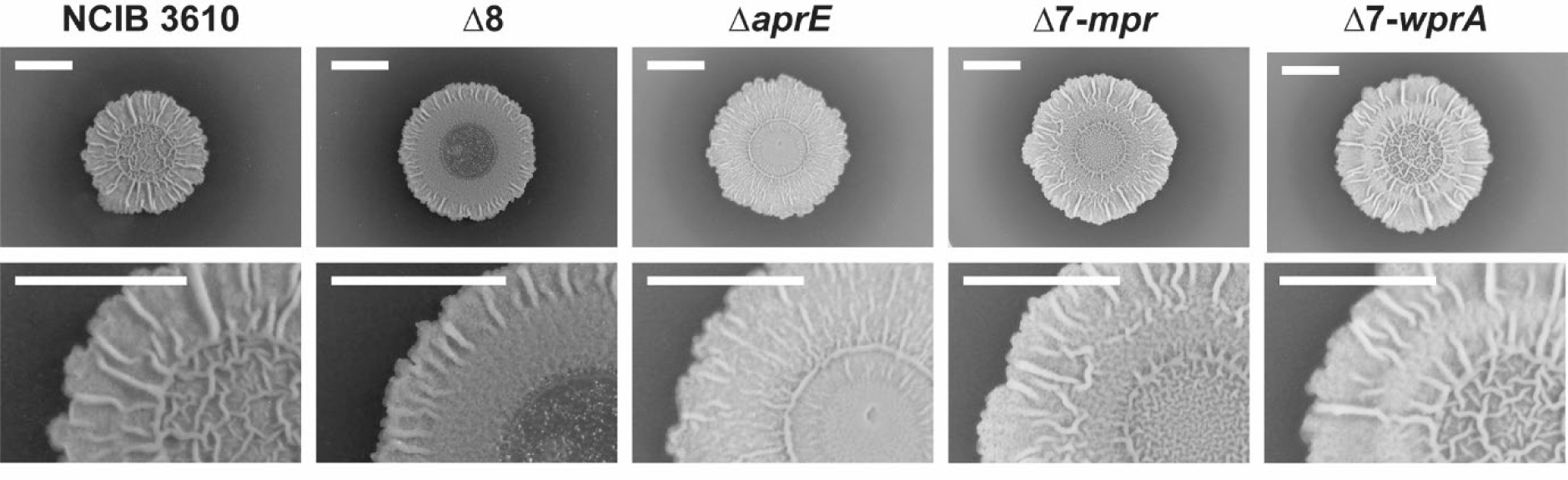
Specific extracellular proteases support the formation of a wrinkled colony biofilm. NCIB 3610, Δ8 and Δ*aprE*, Δ7-*mpr* and Δ7-wprA colony biofilms after 72 hours. All scale bars represent 5 mm. Samples were grown using 0.5% glutamic acid (w/v) and 0.5% glycerol (v/v).

To isolate the role of each extracellular protease and determine its contribution to supporting colony biofilm formation, we repeated the analysis using a library of extracellular protease mono-producer strains (where a single coding region was reinserted into the Δ8 strain (Table 1) (Rosazza et al., 2023)). We observed that only the WprA mono-producer strain had a wrinkle phenotype that was comparable to NCIB 3610 (Figure 4). The Mpr mono-producer strain had a phenotype that was intermediate to that of NCIB 3610 and the Δ8 strain (Figure 4). All the other mono-producer strains only displayed wrinkles at the edge of the biofilm like the Δ8 strain (Figure S3). Quantification of the biofilm footprint area of these strains revealed that producing one of Bpr, Epr, Mpr and WprA increased the ability of *B. subtilis* to expand during biofilm formation (Figure S3), with the footprint occupied increasing beyond that attained by the Δ8 strain. By contrast, restoration of Vpr production to the Δ8 strain impeded the ability of the isolate to expand such that the colony biofilm footprint of the *vpr* monoproducer strain was the same as the WT (Figure S3). The reintroduction of *aprE* or *nrpB* or *nprE* had no impact on the colony biofilm footprint occupied compared with the Δ8 strain (Figure S3). Taken together, these results allowed us to identify that the removal of AprE from the NCIB 3610 secretome negatively impacts colony biofilm architecture and WprA activity alone (where production is driven by its native promoter) is sufficient to promote wrinkle development within the colony biofilm. Moreover, altering the profile of extracellular proteases produced impacts the ability of *B. subtilis* to expand during biofilm formation in a protease-specific manner and a manner that is independent of the ability of the strain to form rugose colony biofilms.

### Reciprocal sharing of two classes of public goods recovers *B. subtilis* colony biofilm morphology

The biofilm matrix and extracellular protease each play an important role in supporting biofilm structure, albeit for different reasons. Mutant strains of each family of public good each have impaired colony biofilm formation phenotypes. Therefore, we hypothesised that reciprocal sharing of the extracellular proteases and the biofilm matrix molecules during co-culture could reinstate a WT NCIB 3610-like phenotype (Figure 5A), as, for example, is the case for the *eps* and *tasA* mutants (Dragos et al., 2018a, Dragos et al., 2018b). To test this, we spotted a mix at an initial 1:1 ratio of either the WT or Δ8 strain expressing mKate with either a single matrix mutant (Δ*bslA,* Δ*eps(A-O)* or Δ*tasA*) or a triple matrix mutant (Δ*bslA,* Δ*eps(A-O),* Δ*tasA*) strain expressing GFP (see Table 1). We evaluated the ability of the co-cultured strains to form a structured biofilm and assessed if the phenotype was comparable to that of the NCIB 3610 strain grown alone under the same conditions. We assessed the proportion of the two strains at the end of the experiments using image analysis methods to determine if the ratio had deviated from the initial 1:1 founder ratio. The image analysis approach was previously extensively evaluated by direct comparison with flow cytometry of the same samples (Eigentler et al., 2022) (see materials and methods for more information). We also quantified the footprint occupied by the biomass. A summary of the findings is provided in Table 2.

**Figure 5.**
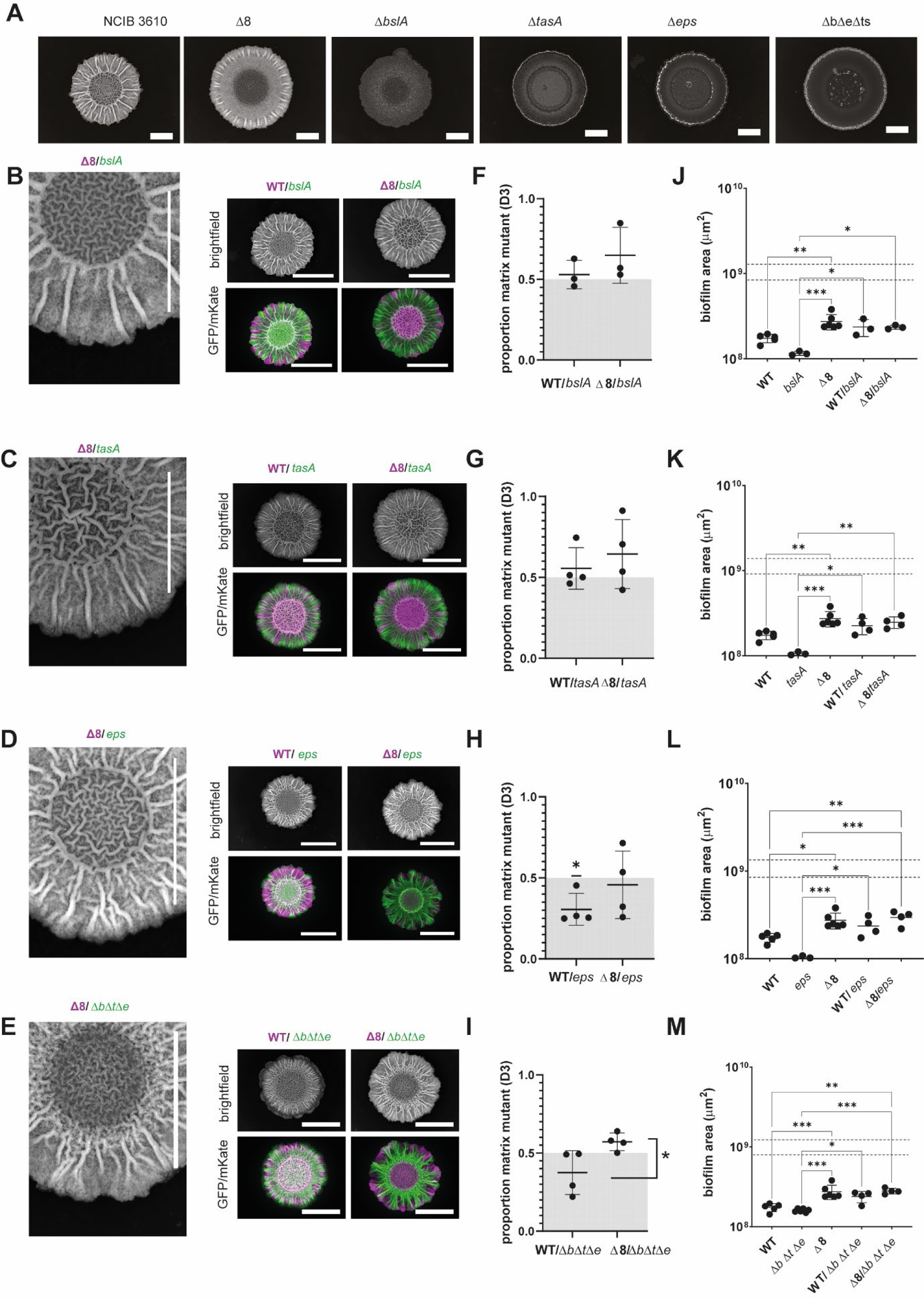
Reciprocal sharing of the extracellular proteases and extracellular matrix molecules restores biofilm formation using glutamic acid as the nitrogen source. **A.** NCIB 3610-mKate, Δ8-mKate, Δ*bslA*, *Δeps(A-O)*, Δ*tasA* and *ΔbslA, Δeps(A-O),ΔtasA* (ΔbΔeΔts) colony biofilms at day 3; **B-E.** Colony biofilms founded using a 1:1 ratio of Δ8-mKate:Δ*bslA*-GFP, Δ8-mKate:Δ*eps(A-O)-*GFP, Δ8-mKate:Δ*tasA*-GFP, or Δ8-mKate:Δ*bslA*, Δ*eps(A-O),*Δ*tasA* (ΔbΔeΔts)-GFP strains at day 3. The raw intensity signal for brightfield is shown in greyscale and the signal from GFP and mKate proteins is false coloured (green=GFP, magenta=mKate); **F-I.** Quantification of the relative density of the GFP and mKate signal within the biofilm footprint. Each point represents the relative density value (n=3-5) and the line represents the mean value and the errors the standard deviation; The deviation of the data points from a theoretical value of 0.5 representing an exact 1:1 ratio of the two starting strains was tested, a statistically significant deviation is indicated by a ‘*’; To assess if there was a statistically significant difference between the outcome driven by the extracellular proteases a standard T-test was used. Statistically significant differences are indicated by a ‘*’; **J-M.** Biofilm footprint area in μm^2^ of NCIB 3610, Δ8 and the cocultures presented in B-E. Each point represents the footprint area value (n=3-5 biological replicates) and the line represents the mean value and the errors the standard deviation. One-way ANOVA was performed between NCIB 3610 and Δ8 or coculture mixes: *** = p-value < 0.001, ** = p-value < 0.01. The scale bar represents 10 mm. A representative image of the independent experiments is shown. Samples were grown using 0.5% glutamic acid (w/v) and 0.5% glycerol (v/v).

**Table 2.** Summary of the reciprocal sharing experiments.

|  | Growth Condition / Feature Assessed | glutamic acid |  | milk |  | BSA |  |
| --- | --- | --- | --- | --- | --- | --- | --- |
| | | NCIB 3610 | $\Delta 8$ | NCIB 3610 | $\Delta 8$ | NCIB 3610 | $\Delta 8$ |
| ***ΔbsIA*** | WT-like structure: | YES | YES | YES | YES | YES | NO (non-wrinkled flat) |
|  | Coexistence: | YES | YES | YES | YES | NO NCIB 3610 wins | YES |
|  | WT-like footprint: | YES | YES | YES | YES | YES | YES |
| ***Δeps(A-O)*** | WT-like structure: | YES | YES | YES | YES | NO (non-wrinkled flat) | NO (non-wrinkled flat) |
| | Coexistence: | NO WT wins | YES | YES | NO $\Delta 8$ wins | NO *Δeps(A-O)* wins | NO *Δeps(A-O)* wins |
|  | WT-like footprint: | YES | NO >WT | YES | YES | YES | YES |
| ***ΔtasA*** | WT-like structure: | YES | YES | NO (irregular edges) | YES | NO (non-wrinkled flat) | NO (non-wrinkled flat) |
|  | Coexistence: | YES | YES | YES | YES | NO NCIB 3610 wins | YES |
|  | WT-like footprint: | YES | YES | YES | YES | YES | YES |
| ***ΔbsIA Δeps(A-O) ΔtasA*** | WT-like structure: | YES | YES | YES | YES | NO (high spatial segregation) | NO (large, flat population) |
|  | Coexistence: | YES | NO matrix mutant wins | YES | NO matrix mutant wins | NO matrix mutant wins | NO matrix mutant wins |
|  | WT-like footprint: | YES | NO > WT | YES | YES | YES | YES |

We observed that compared to the extracellular protease and extracellular matrix component mutants cultured in isolation (Figure 5A), co-culturing the Δ8 strain with any of the single matrix mutants (Δ*bslA,* Δ*eps(A-O)* or Δ*tasA*) or the triple matrix mutant (Δ*bslA,* Δ*eps(A-O),* Δ*tasA*) strain restored NCIB 3610-like colony biofilm formation (compare Figure 5A with Figure 5B, 5C, 5D, 5E). Compared to the flat phenotype of the extracellular matrix component mutant strains, and the phenotype where only the edge of the colony biofilm displayed wrinkles in the case of the Δ8 strain, the presence of both mutant strains in the founding population allowed the formation of wrinkles across the biofilm. The same outcome was observed when we combined the NCIB 3610 strain expressing mKate with any of the single matrix mutants (Δ*bslA,* Δ*eps(A-O)* or Δ*tasA*) or the triple matrix mutant (Δ*bslA,* Δ*eps(A-O),* Δ*tasA*) strains expressing GFP at a 1:1 starting ratio (Figure 5B, 5C, 5D, 5E).

For all combinations tested, the coexistence of the two strains occurred. We quantified the relative density of each population within the mature biofilms using the fluorescent protein reporters and observed that both the Δ8 and NCIB 3610 strains coexisted with the *bslA* and *tasA* matrix mutants, with either the NCBI 3610 or Δ8 strain being present at a similar proportion to the matrix mutants at the end of the experiment (Figure 5F, 5G). When the *eps* mutant was cocultured with the WT strain, the WT showed a slight dominance in the mature biomass (Figure 5H) that was not present when the *eps* mutant was cocultured with the Δ8 strain. Additionally, in the case of the Δ8/Δ*bslA*Δ*tasA*Δ*eps* strain combination, the triple deletion matrix mutant showed an advantage and was quantified as being at a higher proportion compared to when the WT strain was the partner strain (Figure 5I).

All strain combinations exhibited a biofilm footprint area comparable to WT except for the Δ8/Δ*eps* and Δ8/Δ*bslA*Δ*tasA*Δ*eps* strain pairs where the footprint was larger than WT and not statistically different from the Δ8 single strain (Figure 5J, 5K, 5L, 5M). Taken together, we can conclude that reciprocal sharing of the public goods between the Δ8 and matrix mutant strains rescues the WT-like colony biofilm phenotype in these readily accessible nutrient conditions.

### The nutrient source influences biofilm structure and hydrophobicity

To explore the impact on the reciprocal sharing of extracellular proteases and the biofilm matrix molecules in a range of conditions, we increased the pressure placed on the extracellular proteases’ role to sustain growth by adding polymeric nutrient sources. We first needed to identify conditions where biofilm formation occurred, and growth depended on extracellular protease activity. In planktonic conditions, the presence of polymeric nutrient sources forces a dependency on extracellular proteases for growth (Rosazza et al., 2023). Therefore, we removed glutamic acid from the growth medium and replaced it with either 1.5% (w/v) milk or 1% (w/v) bovine serum albumin (BSA). We assessed the ability of NCIB 3610 and the Δ8 strains to form colony biofilms. For the milk-containing medium, rapid growth of a structured biofilm for the NCIB 3610 strain was achieved and, while the Δ8 strain managed to grow, albeit to a lesser extent than the NCIB 3610 strain, it did not manage to form a structured biofilm (Figure 6A) and the footprint the biomass occupied was significantly smaller (Figure 6B). When 1% (w/v) BSA formed the polymeric source of nitrogen, we detected the formation of a colony biofilm for the NCIB 3610 strain. After extended incubation, the colony biofilm showed signs of rugosity, which is indicative of the production of biofilm matrix molecules. The wrinkling was of a different nature than when glutamic acid or milk was used as the nitrogen source (compare with Figure 1B). In the same conditions, growth of the Δ8 strain on the agar surface was significantly impeded (Figure 6A). The Δ8 biomass that was present on the agar surface occupied a smaller footprint than the protease-producing strain (Figure 6B) and showed no evidence of architecture (Figure 6A).

**Figure 6.**
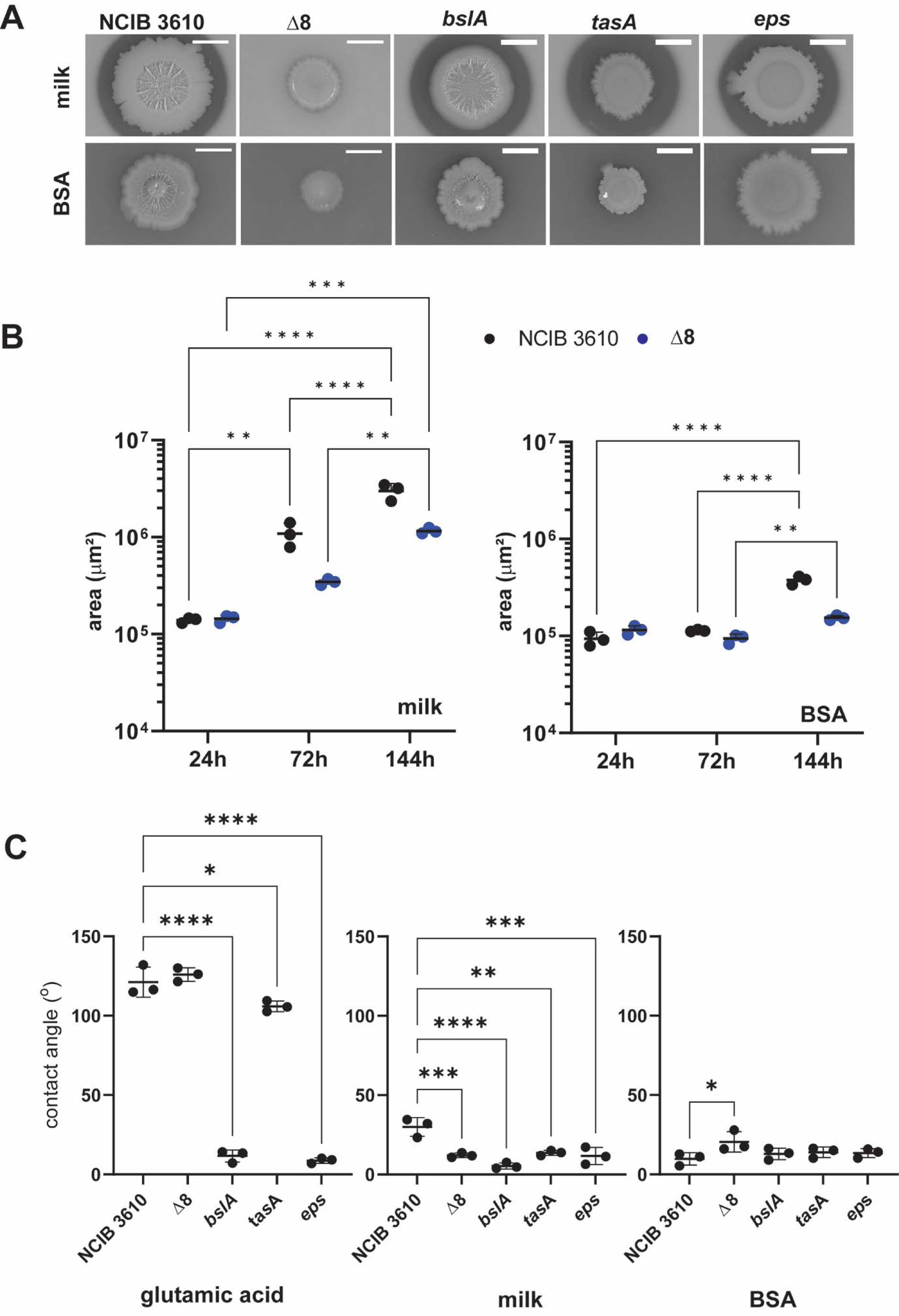
Nutrient source changes biofilm morphology. **A.** NCIB 3610, Δ8 Δ*bslA*, Δ*eps(A-O)* and Δ*tasA* strains colony biofilms at day 3 on 1.5% milk (w/v)) and day 6 (1% BSA (w/v)). The scale bar represents 5 mm. A representative image of three independent experiments is shown. **B.** Biofilm footprint area in *μm^2^* of NCIB 3610, Δ8 and the conditions presented in A. Each point represents the footprint area value (n=3 biological replicates) and the line represents the mean value with the bars being the standard deviation. 2-way ANOVA was used to analyse the data; **C.** Contact angle quantification of a water droplet placed on the surface of NCIB 3610, Δ8 *ΔbslA, Δeps(A-O)* and *ΔtasA* colony biofilms at day 3 (0.5% glutamic acid (w/v) and 1.5% milk (w/v)) and day 6 (1% BSA (w/v)). Each point represents the contact angle values (n=3 biological replicates) and the line represents the mean value.

We did not see a complete lack of growth of the Δ8 strain in either condition. In the medium containing milk, there are likely nitrogen and carbon sources accessible to the Δ8 strain as it shows limited growth even in the absence of extracellular proteases. The same is likely to be true of the medium containing BSA where it is probable that small peptides are present that facilitate the limited growth observed by the Δ8 strain. Over time, it is also likely that some cells in the biomass will sporulate or lyse thereby releasing intracellular proteases that allow the degradation of exogenous proteins allowing the Δ8 to show limited growth. However, we concluded that there was a sufficiently distinct dependence on the extracellular proteases for growth to progress.

We assessed the properties of the colony biofilms formed under these conditions and tested their dependency on the known extracellular matrix molecules to support colony biofilm structure. When the nutrient source was milk, we observed that each matrix mutant strain (Δ*bslA,* Δ*eps,* or Δ*tasA*) had a halo surrounding them where the milk contained in the substratum had been degraded (Figure 6A). Thus, the extracellular matrix mutants produce extracellular proteases. The Δ*eps(A-O)* and Δ*tasA* mutants grew on all these media and, as noted for other conditions and contexts (Dragos et al., 2018a), had a defect in forming a structured colony biofilm (Figure 6A). In contrast to glutamic acid driven growth, the Δ*bslA* strain still formed a structured biofilm on both milk and BSA-containing growth media. When grown using glutamic acid as the nitrogen source, NCIB 3610 colony biofilms have a non-wetting surface. In contrast, the contact angle of the water droplet placed on NCIB 3610 biofilm grown on either polymeric nutrient source media was below 90° (indicating a hydrophilic or wetting surface) (Figure 6C). The contact angle findings would be consistent with BslA not being produced, however, we detected BslA (and TasA) after immunoblotting of protein extracts obtained from the NCIB 3610 biofilm grown using milk or BSA as the polymeric nitrogen source (Figure S3). These results indicate that BslA is produced, but largely dispensable for biofilm structure in these nutrient conditions, consistent with its reported dispensability for biofilm formation in some conditions, for example on fungal hyphae (Kjeldgaard et al., 2019).

### Reciprocal sharing of public goods becomes restricted as the dependency on the extracellular proteases for growth rises

Having established conditions for *B. subtilis* colony biofilm formation where growth depends on the extracellular proteases, we next assessed the impact of co-culture when we combined the protease mutant with extracellular matrix mutants. We used the same process as before and a summary of the findings is provided in Table 2.

When milk provided the nitrogen source, a recovery towards the morphology of the NCIB 3610 phenotype and area occupied on the same conditions was observed when the Δ8 strain was co-cultured with any of the *bslA*, *eps(A-O)*, *tasA* single mutant strains or the triple matrix mutant (Figures 7A-7E). The inclusion of the matrix mutant in coculture with the WT strain did not impact colony biofilm phenotype or footprint, except for the *tasA* strain where the colonies appeared more irregular (Figure 7C). We established that the combination of strains coexists in all tested pairs. However, for the Δ8:*eps(A-O)* strain combination the Δ8 strain displayed increased abundance over the *eps(A-O)* mutant (Figure 7H) and for the Δ8/Δ*bslA*Δ*tasA*Δ*eps* mutant combination, the Δ*bslA*Δ*tasA*Δ*eps* strain dominated (Figure 7I). Collectively, these data show that genetically distinct isolates can share the two classes of public goods to allow growth and enhanced biofilm formation (Figure 7J-M). Moreover, we established that the triple matrix mutant strain can take advantage of the Δ8 strain but not the WT strain to enhance its ability to occupy a surface and form a biofilm (Figure 7E, I, M).

**Figure 7.**
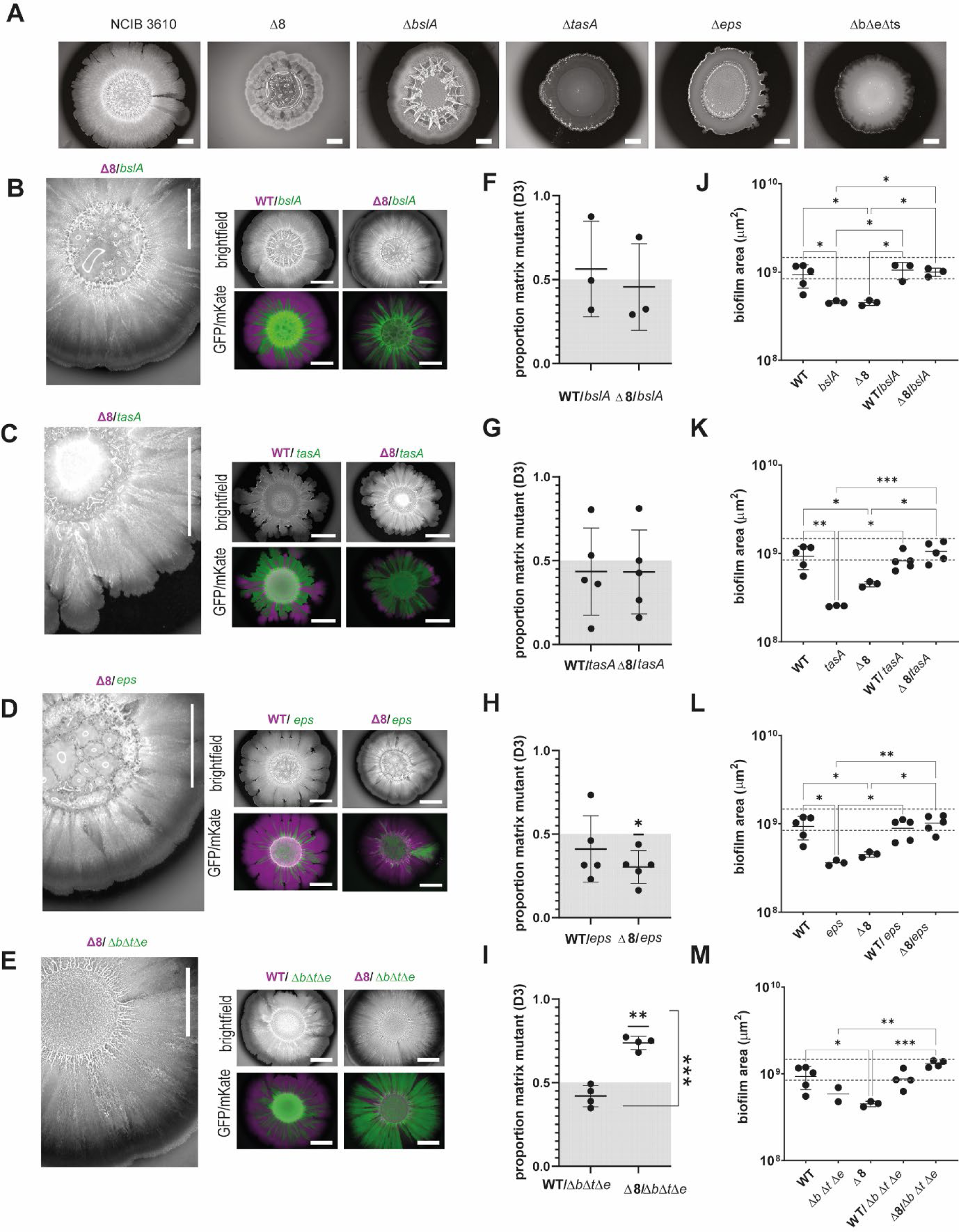
Reciprocal sharing of the extracellular proteases and extracellular matrix molecules restores biofilm formation using milk as the nitrogen source. **A.** NCIB 3610-mKate, Δ8-mKate, Δ*bslA*, *Δeps(A-O)*, Δ*tasA* and *ΔbslA, Δeps(A-O),ΔtasA* (ΔbΔeΔts) colony biofilms at day 3; **B-E.** Colony biofilms founded using a 1:1 ratio of Δ8-mKate:Δ*bslA*-GFP, Δ8-mKate:Δ*eps(A-O)-*GFP, Δ8-mKate:Δ*tasA*-GFP, or Δ8-mKate:Δ*bslA*, Δ*eps(A-O),*Δ*tasA* (ΔbΔeΔts)-GFP strains at day 3. The raw intensity signal for brightfield is shown in greyscale and the signal from GFP and mKate proteins is false coloured (green=GFP, magenta=mKate); **F-I.** Quantification of the relative density of the GFP and mKate signal within the biofilm footprint. Each point represents the relative density value (n=3-5) and the line represents the mean value and the errors the standard deviation; The deviation of the data points from a theoretical value of 0.5 representing an exact 1:1 ratio of the two starting strains was tested, a statistically significant deviation is indicated by a ‘*’; To assess if there was a statistically significant difference between the outcome driven by the extracellular proteases a standard T-test was used. Statistically significant differences are indicated by a ‘*’; **J-M.** Biofilm footprint area in μm^2^ of NCIB 3610, Δ8 and the cocultures presented in B-E. Each point represents the footprint area value (n=3-5 biological replicates) and the line represents the mean value and the errors the standard deviation. One-way ANOVA was performed between NCIB 3610 and Δ8 or coculture mixes: *** = p-value < 0.001, ** = p-value < 0.01; ** = p-value < 0.05. The scale bar represents 10 mm. A representative image of the independent experiments is shown. Samples were grown using 0.5% glutamic acid (w/v) and 0.5% glycerol (v/v).

When the dependence on the extracellular proteases for growth was further increased (as evident from the reduced biofilm footprint (Figure 6)), which was achieved by replacing milk with BSA, the ability of the two founding strains to recover a wild-type-like biofilm morphology and to coexist decreased (Figures 8A-8E). Analysis of the population composition (Figure 8F-I) indicated that the Δ8 strain had a competitive advantage when co-cultured with either the single *bslA* or *tasA* mutants compared with the WT strain (Figure 8F-G). In contrast, both the WT and Δ8 had a lower competitive advantage when mixed with the *eps(A-O)* mutant or the triple Δ*bslA*Δ*tasA*Δ*eps* strains (Figure 8H-I). In these cases, the matrix mutant dominated the mature population suggesting that not producing the exopolysaccharide provides a significant growth advantage from which neither the Δ8 nor WT strain benefits (Figure 8J-M).

**Figure 8.**
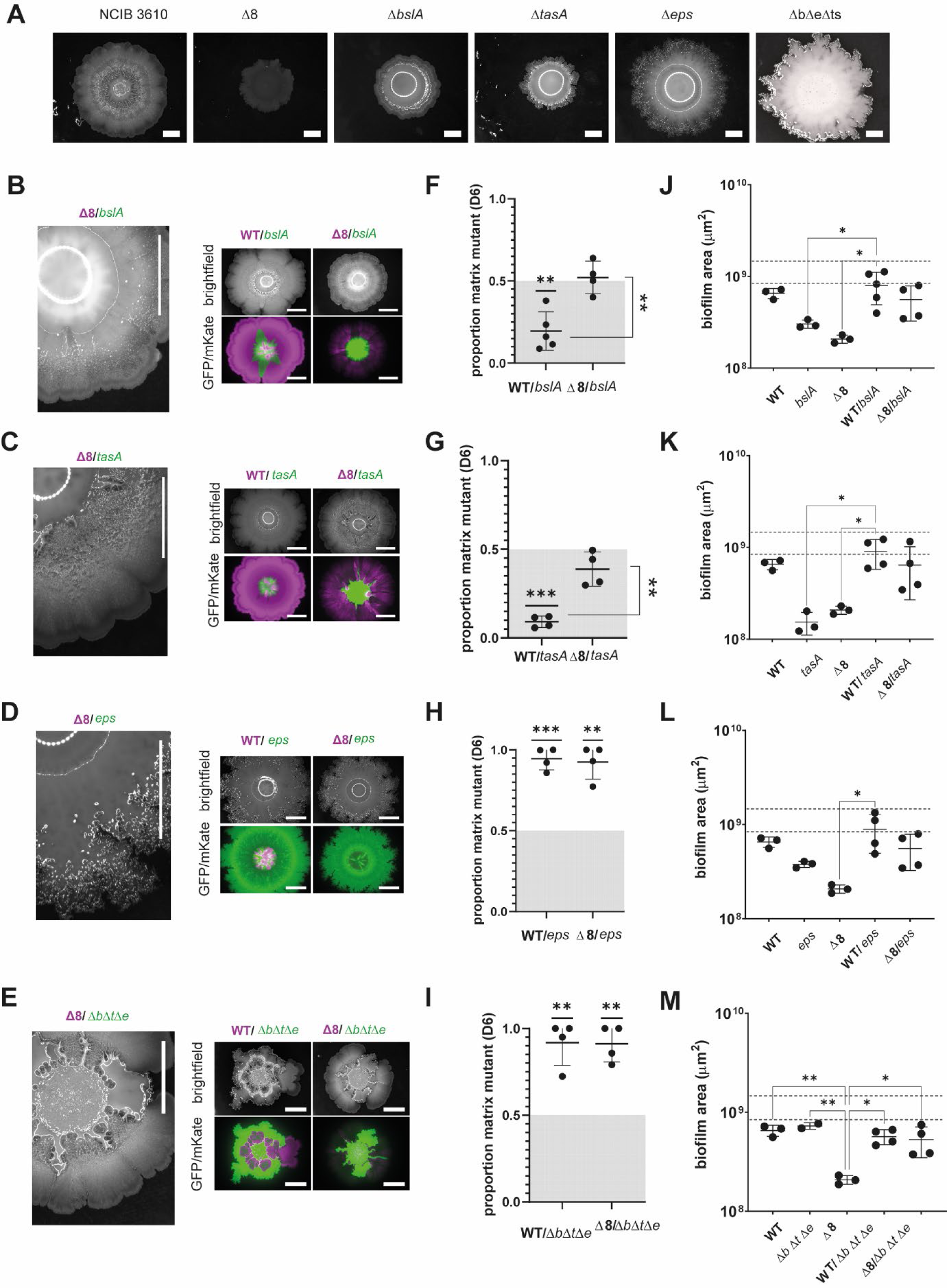
Reciprocal sharing of the extracellular proteases and extracellular matrix molecules restores biofilm formation using BSA as the nitrogen source. **A.** NCIB 3610-mKate, Δ8-mKate, Δ*bslA*, *Δeps(A-O)*, Δ*tasA* and *ΔbslA, Δeps(A-O),ΔtasA* (ΔbΔeΔts) colony biofilms at day 3; **B-E.** Colony biofilms founded using a 1:1 ratio of Δ8-mKate:Δ*bslA*-GFP, Δ8-mKate:Δ*eps(A-O)-*GFP, Δ8-mKate:Δ*tasA*-GFP, or Δ8-mKate:Δ*bslA*, Δ*eps(A-O),*Δ*tasA* (ΔbΔeΔts)-GFP strains at day 3. The raw intensity signal for brightfield is shown in greyscale and the signal from GFP and mKate proteins is false coloured (green=GFP, magenta=mKate); **F-I.** Quantification of the relative density of the GFP and mKate signal within the biofilm footprint. Each point represents the relative density value (n=3-5) and the line represents the mean value and the errors the standard deviation; The deviation of the data points from a theoretical value of 0.5 representing an exact 1:1 ratio of the two starting strains was tested, a statistically significant deviation is indicated by ‘**’ or ‘***’; To assess if there was a statistically significant difference between the outcome driven by the extracellular proteases a standard T-test was used. Statistically significant differences are indicated by a ‘*’; **J-M.** Biofilm footprint area in *μm^2^* of NCIB 3610, Δ8 and the cocultures presented in B-E. Each point represents the footprint area value (n=3-5 biological replicates) and the line represents the mean value and the errors the standard deviation. One-way ANOVA was performed between NCIB 3610 and Δ8 or coculture mixes: *** = p-value < 0.001, ** = p-value < 0.01; * = p-value < 0.05. The scale bar represents 10 mm. A representative image of the independent experiments is shown. Samples were grown using 0.5% glutamic acid (w/v) and 0.5% glycerol (v/v).

## Discussion

Public goods are essential in microbial communities and are diverse in form and function (Momeni, 2018). Public goods are often produced by a subset of cells within the biofilm and the resultant materials are shared with other cells, even those not directly involved in production (Bachmann et al., 2013, Martin et al., 2020). This results in the potential for exploitation by non-producers. *B. subtilis* biofilms contain multiple public goods and, in this context, are typically thought of as the extracellular matrix molecules (Arnaouteli et al., 2021). The extracellular matrix molecules are shared to different degrees in cell populations with some being restricted more closely to the producing cells (Jautzus et al., 2022). The results presented here provide insight into the role of extracellular proteases, a newly defined biofilm public good, in the formation of *B. subtilis* colony biofilms. The extracellular proteases play an important role in biofilm formation, even when nitrogen is in the form of glutamic acid. These are nutrient conditions where growth under planktonic conditions is not impacted by their absence (Rosazza et al., 2023), thereby showcasing the impact of different environmental conditions. It is possible that materials released from dead cells in the biofilm (Asally et al., 2012) or from re-modelling / degradation of protein components in the matrix (for example the protein TapA which is required for biofilm matrix formation, (Earl et al., 2020)) are not accessible when these lytic enzymes are absent and therefore nutrients become limited in localised regions resulting in increased numbers of spores.

The role of individual extracellular proteases produced by *B. subtilis* in promoting biofilm formation was systematically investigated using a suite of single mutant and single producer strains derived from NCIB 3610. Interestingly, most single deletion strains did not significantly affect colony biofilm phenotype, with the architecture of the colony biofilms that formed mirroring the wild-type strain and with the findings for Vpr and NprE being consistent with previous work (Earl et al., 2020). However, the absence of AprE resulted in a distinct phenotype with smaller, scattered wrinkles, revealing a new role for AprE in colony biofilm formation. Furthermore, a WprA mono-producer strain (and to a lesser extend the Mpr monoproducer strain) exhibited a similar wrinkle phenotype to NCIB 3610, while other mono-producer strains resembled the Δ8 strain. These findings indicate that WprA is sufficient to allow wild-type-like biofilm architecture, albeit the targets are unknown and may include TasA. Collectively, these observations suggest that specific proteases have unique functions in biofilm formation, although for the most part, their target(s) remain undefined (Harwood and Kikuchi, 2022).

By modulating the nutrient source, we were able to influence both biofilm structure and hydrophobicity. The presence of polymeric nutrient sources increases the dependency on extracellular proteases for growth (Rosazza et al., 2023). This dependency impacts biofilm formation, particularly under conditions where protease production becomes more costly (Rosazza et al., 2023). The same is true in the case of biofilm formation. The findings highlight how the typical composition of the media used in a field of research can influence the experimental outcomes revealed and may even prevent important insights from being uncovered. Our data provide a platform to expand the conditions in *B. subtilis* biofilm formation can be analysed and showcase the adaptability of *B. subtilis* to varying nutrient sources and reveal the associated trade-offs in terms of protease production and biofilm structure.

We explored the concept of reciprocal sharing of extracellular proteases and matrix components in co-culture experiments and found that co-culturing the Δ8 strain with matrix mutant strains restores wild-type biofilm morphology, indicating that public good sharing can rescue biofilm structure. However, the relative cost of protease production influences coexistence within these communities, as shown by experiments where the dependency on the extracellular proteases for growth is increased. *B. subtilis* produces other classes of extracellular enzymes, such as lipases, and amylases (Eggert et al., 2003, Konsula and Liakopoulou-Kyriakides, 2004). Thus, it is highly likely that these groups of enzymes also serve as public goods that become essential in specific growth contexts.

### Outlook

Our findings provide insights into the intricate interplay between extracellular proteases and the biofilm matrix molecules in the context of public goods and biofilm formation in *B. subtilis*. It also raises intriguing questions about the specific functions of individual extracellular proteases and the ecological implications of public goods sharing within biofilm communities. Understanding public goods sharing in *B. subtilis* biofilms is not only relevant from a biological perspective, but also has practical implications. Increased knowledge should provide insights into biofilm resilience, the development of pro-biofilm strategies, and thus the potential for engineering microbial communities for various applications.

## Acknowledgements

Work was funded by the Biotechnology and Biological Science Research Council (BBSRC) [BB/P001335/1, BB/R012415/1]. TR was funded by Wellcome [102132/B/13/Z]. This research was funded in whole, or in part, by the Wellcome Trust [102132/B/13/Z]. CE was supported by the Wellcome Institutional Strategic Support Fund [Award no. 097818/Z/11]. The authors would like to thank Dr. Sofia Arnaouteli and Ms Tetyana Sukhodub for their support in generating some of the strains used in this study. The authors thank Dr. Megan Bergkessel for her constructive discussions on this study. **For the purpose of open access, the author has applied a CC BY public copyright licence to any Author Accepted Manuscript version arising from this submission.**

## Data Availability Statement

Analysis code and experimental datasets have been deposited in the nrstanleywall GitHub repository (https://github.com/NSWlabDundee/) and archived by Zenodo (Rosazza et al., 2024).

## Supplemental Material

**Figure S1.**
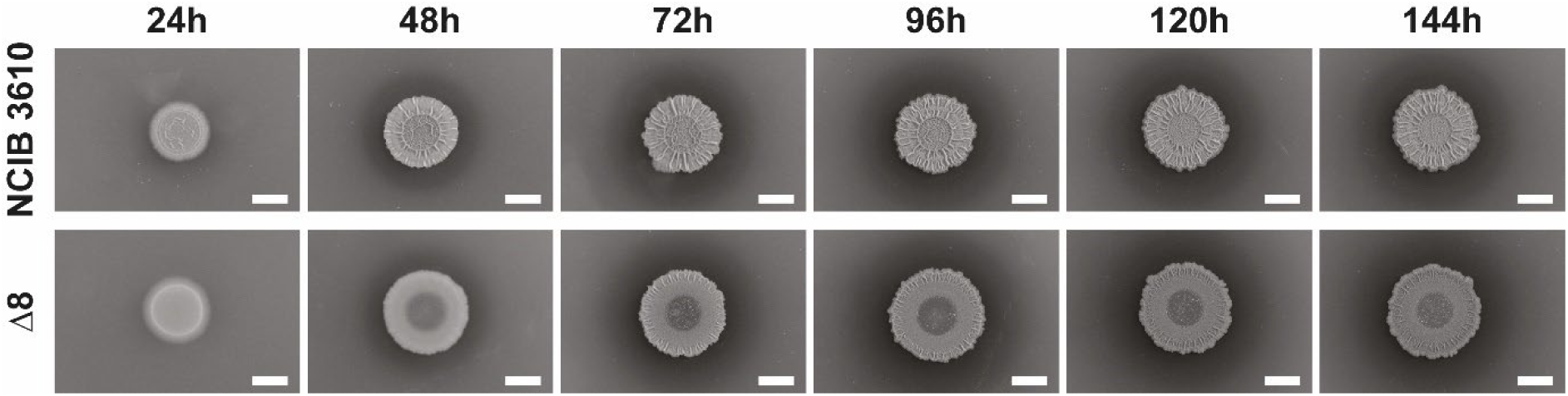
NCIB 3610 and Δ8 colony biofilms grown for up to 144 hours using MS base supplemented with 0.5% glutamic acid (w/v). A representative image of five independent experiments is shown. Scale bar represents 5 mm.

**Figure S2.**
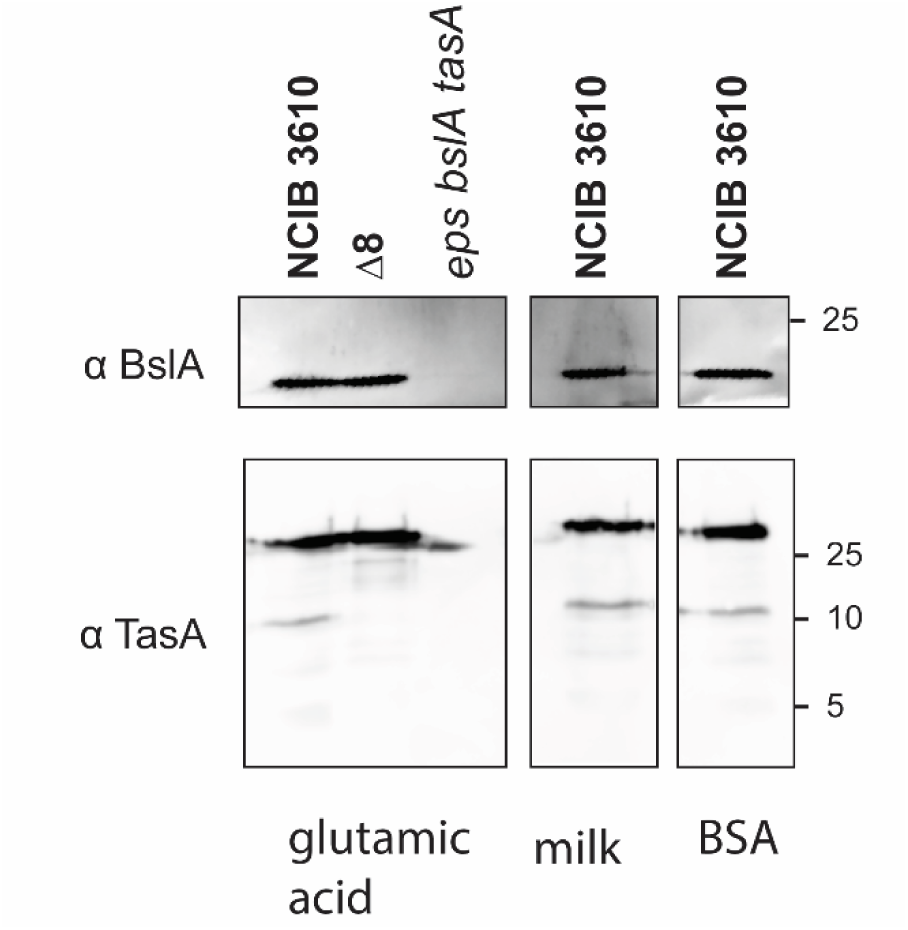
Extracellular matrix proteins are produced in when nutrients are polymeric. Images of immunoblot against BslA (top image) and TasA (bottom image) on biofilm protein extract obtained after 3 days from NCIB 3610, Δ8 and the triple *ΔbslA, Δeps(A-O), ΔtasA* (*eps bslA tasA*) strains. A representative image of three independent experiments is shown (left part of the image correspond to Figure 2).

**Figure S3.**
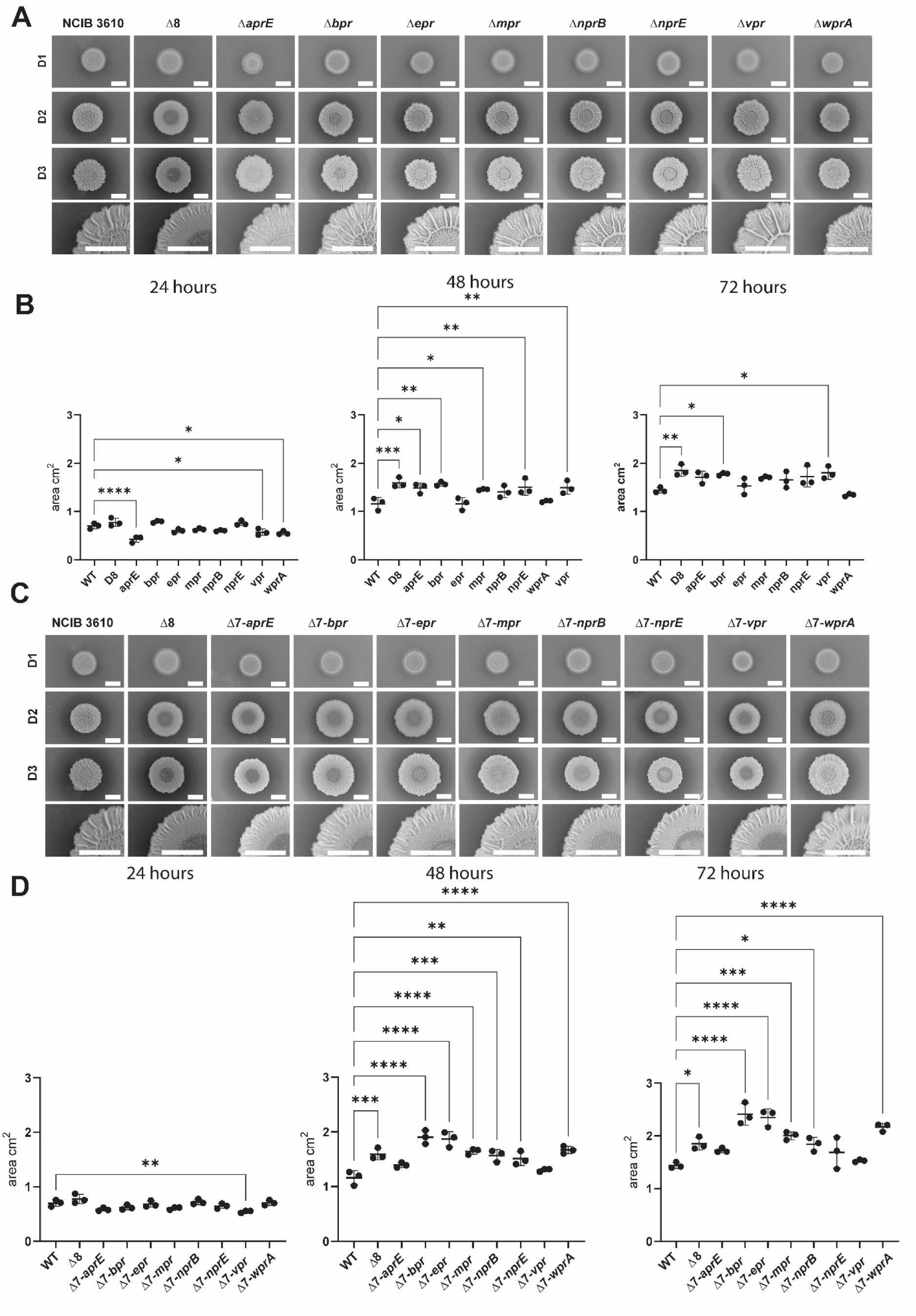
Single deletion and monoproducers strains exhibit different biofilm phenotypes. **A.** Biofilm reflected light images of NCIB 3610, Δ8 and single deletion strains over time (day 1 to day 3). A zoom of the day 3 image is in the bottom row. The scale bar represents 5 mm; **B.** Biofilm footprint area in cm^2^ of NCIB 3610, Δ8 and single deletion strains over time. Each point represents the footprint area value of n=3 biological repeats and the line represents the mean value. **C.** Biofilm reflected light images of NCIB 3610, Δ8 and monoproducer strains over time (day 1 to day 3). Biofilm footprint area in cm^2^ of NCIB 3610, Δ8 and single deletion strains over time. Each point represents the footprint area value of n=3 biological repeats and the line represents the mean value. The scale bar represents 5 mm. The same WT and Δ8 samples are presented on each graph. Ordinary One-way ANOVA with multiple comparisons with respect to the WT strain was performed. Only statistically significant differences are shown. *\*\*\*\* = p-value < 0.0001, *** = p-value < 0.001, ** = p-value < 0.01, * = p-value < 0.05*.

